# Unraveling the molecular basis of host cell receptor usage in SARS-CoV-2 and other human pathogenic β-CoVs

**DOI:** 10.1101/2020.08.21.260745

**Authors:** Camila Pontes, Victoria Ruiz-Serra, Rosalba Lepore, Alfonso Valencia

## Abstract

The recent emergence of the novel SARS-CoV-2 in China and its rapid spread in the human population has led to a public health crisis worldwide. Like in SARS-CoV, horseshoe bats currently represent the most likely candidate animal source for SARS-CoV-2. Yet, the specific mechanisms of cross-species transmission and adaptation to the human host remain unknown. Here we show that the unsupervised analysis of conservation patterns across the β-CoV spike protein family, using sequence information alone, can provide rich information on the molecular basis of the specificity of β-CoVs to different host cell receptors. More precisely, our results indicate that host cell receptor usage is encoded in the amino acid sequences of different CoV spike proteins in the form of a set of specificity determining positions (SDPs). Furthermore, by integrating structural data, *in silico* mutagenesis and coevolution analysis we could elucidate the role of SDPs in mediating ACE2 binding across the Sarbecovirus lineage, either by engaging the receptor through direct intermolecular interactions or by affecting the local environment of the receptor binding motif. Finally, by the analysis of coevolving mutations across a paired MSA we were able to identify key intermolecular contacts occurring at the spike-ACE2 interface. These results show that effective mining of the evolutionary records held in the sequence of the spike protein family can help tracing the molecular mechanisms behind the evolution and host-receptors adaptation of circulating and future novel β-CoVs.

**Significance:** Unraveling the molecular basis for host cell receptor usage among β-CoVs is crucial to our understanding of cross-species transmission, adaptation and for molecular-guided epidemiological monitoring of potential outbreaks. In the present study, we survey the sequence conservation patterns of the β-CoV spike protein family to identify the evolutionary constraints shaping the functional specificity of the protein across the β-CoV lineage. We show that the unsupervised analysis of statistical patterns in a MSA of the spike protein family can help tracing the amino acid space encoding the specificity of β-CoVs to their cognate host cell receptors. We argue that the results obtained in this work can provide a framework for monitoring the evolution of SARS-CoV-2 specificity to the hACE2 receptor, as the virus continues spreading in the human population and differential virulence starts to arise.

## Introduction

The emergence of the novel SARS-CoV-2 and its ability to infect humans underscores the epidemic potential of coronaviruses, with the ongoing outbreak being the third documented event in humans in the last two decades (1, 2). To date (August 2020), there have already been more than 20 million reported cases of COVID-19 worldwide and over 770,000 deaths (https://covid19.who.int/) (3).

Coronaviruses (CoVs) are enveloped, positive-sense RNA viruses known for infecting a wide range of hosts (4). Together with SARS-CoV, MERS-CoV, OC43 and HKU1 from the β-CoV genus, and 229E and NL63 from the α-CoV genus, SARS-CoV-2 is the seventh confirmed CoV able to infect humans. While 229E, OC43, NL63 and HKU1 widely circulate in the human population and mostly cause mild disease manifestations in immunocompetent individuals, SARS and MERS CoVs are mainly spread in zoonotic reservoirs, with different intermediate host putatively involved in human transmission (4). CoV entry into the host cells is mediated by the spike glycoprotein, a membrane-anchored homotrimer consisting of two distinct subunits: S1, responsible of viral-host recognition and S2, promoting virus-cell membrane fusion (5, 6). Similar to SARS-CoV, SARS-CoV-2 S has been reported to bind the human angiotensin converting enzyme 2 (hACE2) as a cellular entry receptor via the receptor binding domain (RBD), located in the C-terminal region of the S1 subunit (7).

Both SARS-CoV-2 and SARS-CoV belong to the Sarbecovirus subgenus, their spike proteins share more than 75% sequence identity and show similar RBD architecture and mode of binding to the human receptor (8). However, structural and biophysical evidence showed that SARS-CoV-2 S exhibits a 10-to 20-fold increased affinity to hACE2 compared to its SARS counterpart (7) and, compared to other β-CoVs, possesses unique features such as a putative O-glycosylation site and a polybasic cleavage site (9). The latters are invoked as key factors to be monitored towards understanding the differential virulence of SARS-CoV-2, while determinants of host adaptation and cellular tropism are mainly found within the S1 subunit (7, 10).

Like in SARS-CoV, horseshoe bats represent the most likely candidate animal source for SARS-CoV-2 so far (11). Yet, the specific mechanisms of cross-species transmission and adaptation to the human host remain unknown. Here, we survey the sequence patterns of the β-CoV spike protein family to elucidate the molecular events involved in the differential host-pathogen interaction patterns observed across β-CoVs. **Our results show that receptor specificity is encoded in the amino acid sequences of different β-CoVs spike proteins in the form of a set of specificity determining positions (SDPs)**. We argue that both the methodology and results presented in this work can help tracing the molecular mechanisms behind the evolution and host-receptors adaptation of circulating and future novel coronaviruses.

## Methods

### Sequence dataset and multiple sequence alignment

A BLAST search was performed against the NCBI nr database using the SARS-CoV-2 spike protein (NCBI Reference Sequence: YP_009724390.1) as a query (12). The top 1000 significant hits (e-value <0.05) were selected and clustered to identify non redundant sequences using CD-Hit (13) with the following parameters: -s 0.90 -c 0.98. Cluster representatives were used to build a multiple sequence alignment (MSA) using the MAFFT software (14). The final alignment contained 135 spike protein sequences which were manually reviewed to annotate information on viral strain, host organism and cell receptors using information extracted from the NCBI Protein database, UniprotKB and the literature.

### Detection of SDPs and spike protein subfamilies

**The S3Det method (15) uses multiple correspondence analysis (MCA) to identify differentially conserved positions and sequence subfamilies within a given MSA. MSA positions that follow the subfamily segregation are defined as specificity determining positions (SDPs) of the family**. Here, the unsupervised mode of S3Det was used in a two-level decomposition analysis to identify SDPs linked to the spike protein family segregation between and within β-CoV subgroups. Phylogenetic analysis was performed both on the full β-CoV MSA and for individual subfamilies using the PhyML method (16) with default parameters.

### Coevolution analysis

Coevolving MSA positions were identified by computing the MI-APC (17). MI-APC is a mutual information (MI)-based score corrected by the average product correction (APC) of the background noise and phylogenetic signal. To ensure robust statistics, MSA columns were filtered according to percentage of gaps (<= 50%) and Shannon entropy (>= avg - std), computed as follows:

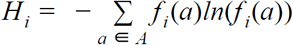

where *Hi* is the Shannon entropy of the *i* -th MSA position, *f _i_* (*a*) is the frequency of amino acid *a* in the *i* -th MSA position, and *A* is the alphabet of all possible amino acids.

### Protein domain annotation and enrichment analysis

Domain enrichment analyses were performed by hypergeometric testing and p-values computed as follows:

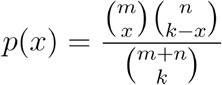

where *x* is the number of SDPs observed in a given domain of interest, *k* is the number of SDPs in the test set, *m* is the length of the domain of interest and *n* + *m* is the size of the protein. The analyses were performed both at the subunit and domain level. Annotations on subunits and domain boundaries of the SARS-CoV-2 spike glycoprotein were retrieved from the literature (18) and mapped to the rest of human β-CoVs spike sequences used in this analysis (SARS-CoV, MERS, OC43 and HKU1) based on the MSA.

### Evolutionary rate analysis

Per-site evolutionary rates (EV) were computed by Rate4Site (19) using as input the full MSA of the β-CoV spike family and phylogenetic tree. SARS-CoV-2 sequence was set as the reference sequence. Raw rate values were used to compute relative rates by normalizing the values to the mean of 1 (20).

### Structural dataset

All the structures employed in this study were retrieved from the PDB (https://www.rcsb.org) using the following PDB codes: 6VSB (7), 6VXX (21), 6LZG (22), 5×5B (23), 5×58 (23), 2AJF (24), 6OHW (25), 6NZK (25), 5×5F (23), 4L72 (26) and 5I08 (27).

## Results

### SDPs as predictors of receptor specificity across the β-CoV lineage

Here we aim at analysing the effect of evolutionary constraints in shaping the organisation of the spike protein family across the β-CoV lineage. To this aim, we use a multivariate-based protocol which allows the automatic and simultaneous detection of both the protein family segregation and associated amino acid variations, i.e. SDPs.

A phylogenetic analysis was performed based on a MSA of the full-lenght spike sequences from SARS-CoV-2 and other representative β-CoVs. The resulting tree, shown in Figure 1A, reflects the taxonomic classification of β-CoVs into five subgenera, namely Sarbecovirus, Hibecovirus, Nobecovirus, Embecovirus and Merbecovirus (29). A first S3Det analysis was performed on individual protein subfamilies from the Sarbecovirus, Embecovirus and Merbecovirus subgroups. In the case of Sarbecovirus, we identified three different clusters, two of which correspond to known Sarbecovirus clades (30). Notably, in contrast to what observed based on phylogenetic analysis, both SARS-CoV-2 and RaTG13 are clustered together with members of Sarbecovirus clade 1, which includes human SARS-CoV and bat SARS-like sequences (Figure 1B-C). This result is confirmed by the analysis of similarity scoring matrices (Supplementary Figure S1) where SARS-CoV-2 and SARS-CoV sequences cluster together and are closer to the green cluster based on the similarity of SDPs, but not when the full-length sequence is considered. Within the Embecovirus group, we identified three clusters: a first cluster corresponding to murine CoVs (MHV), a second containing rat (RtCoVs) and human CoVs (CoV-HKU1), and a third cluster containing the human CoV hCoV-OC43 and other mammalian CoVs (Supplementary Figure S2-A,C). Within the Merbecovirus, we identified four clusters: a first cluster containing MERS and MERS-like bat CoVs, a second cluster containing hedgehog and bat CoVs, a third cluster containing HKU5 bat CoVs, and a fourth cluster containing one single bat CoV isolate (KW2E-F93) from the Nycteria species (Supplementary Figure S2-B,D).

**Figure 1.**
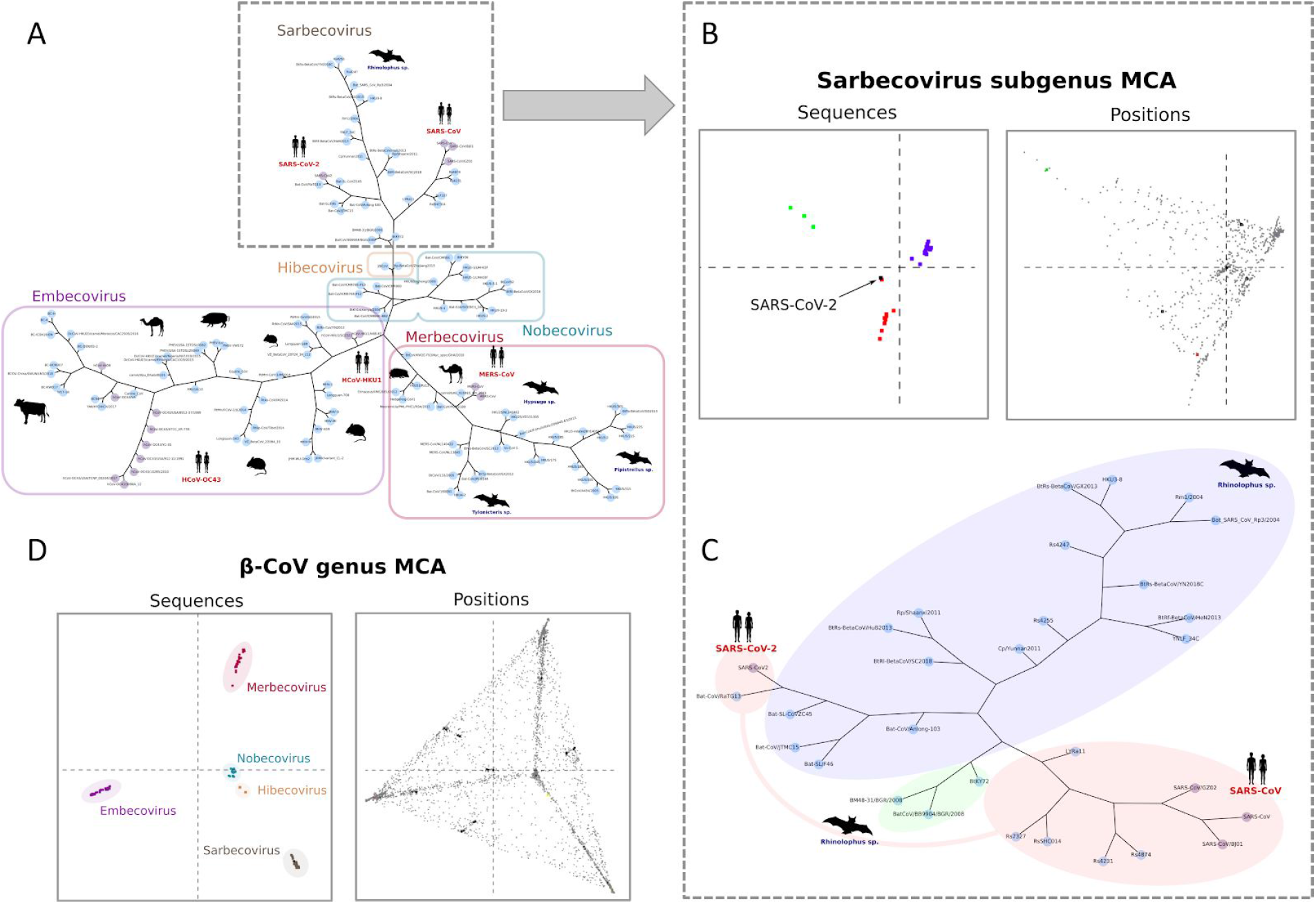
A) Phylogenetic tree obtained for the complete β-CoV spike protein family. S3Det subfamilies are highlighted by rectangles and reflect the phylogenetic classification of Betacoronavirus into five subgenera. B) Results of the S3Det MCA analysis showing the subfamily segregation and associated amino acid positions obtained for the Sarbecovirus subgroup. C) Phylogenetic tree obtained for the Sarbecovirus subgroup. S3Det clusters are highlighted by red, green and blue silhouettes. Both SARS-CoV-2 and RaTG13 are clustered together with SARS-CoV and other members of Sarbecovirus clade 1. D) Results of the S3Det MCA analysis showing the subfamily segregation and associated amino acid positions obtained for the full β-CoV family. Phylogenetic trees were built using PhyML (28). Nodes representing spike protein sequences from human pathogenic CoVs are colored in purple. Host species are shown for some of the nodes as dark silhouettes.

The SDPs associated with the subfamily segregation within these three subgenera display a strong domain enrichment within the spike S1 subunit (Figure 2, Supplementary Table S1). Specifically, the SDPs (n = 28) of Sarbecovirus subfamilies, containing both SARS-CoV and SARS-CoV-2 spike sequences, are enriched in the RBD whereas in hCoV-OC43 and hCoV-HKU1, the human-pathogenic species belonging to Embecovirus, SDPs (n = 12) fall mainly in an upstream region of the spike, with a significant enrichment in the NTD. Also in MERS-CoV we observe a significant enrichment within the S1 subunit. However, as shown in Figure 2, the SDPs (n = 33) are almost evenly distributed across the NTD and RBD regions, with a significant enrichment in the latter (Supplementary Table S2). **Notably, the distribution of the SDPs shows a clear relationship with the cell receptor usage observed among β-CoVs**. In particular, hCoV-HKU1 and hCoV-OC43 are known to bind sialic acid receptors on the host cells via the spike NTD (31), SARS-CoV and SARS-CoV-2 recognize the ACE2 receptors via the RBD (32, 33), while MERS-CoV has been reported to use both DPP4 (34) and sialic acid receptors (35) via the RBD and NTD domains, respectively.

**Figure 2.**
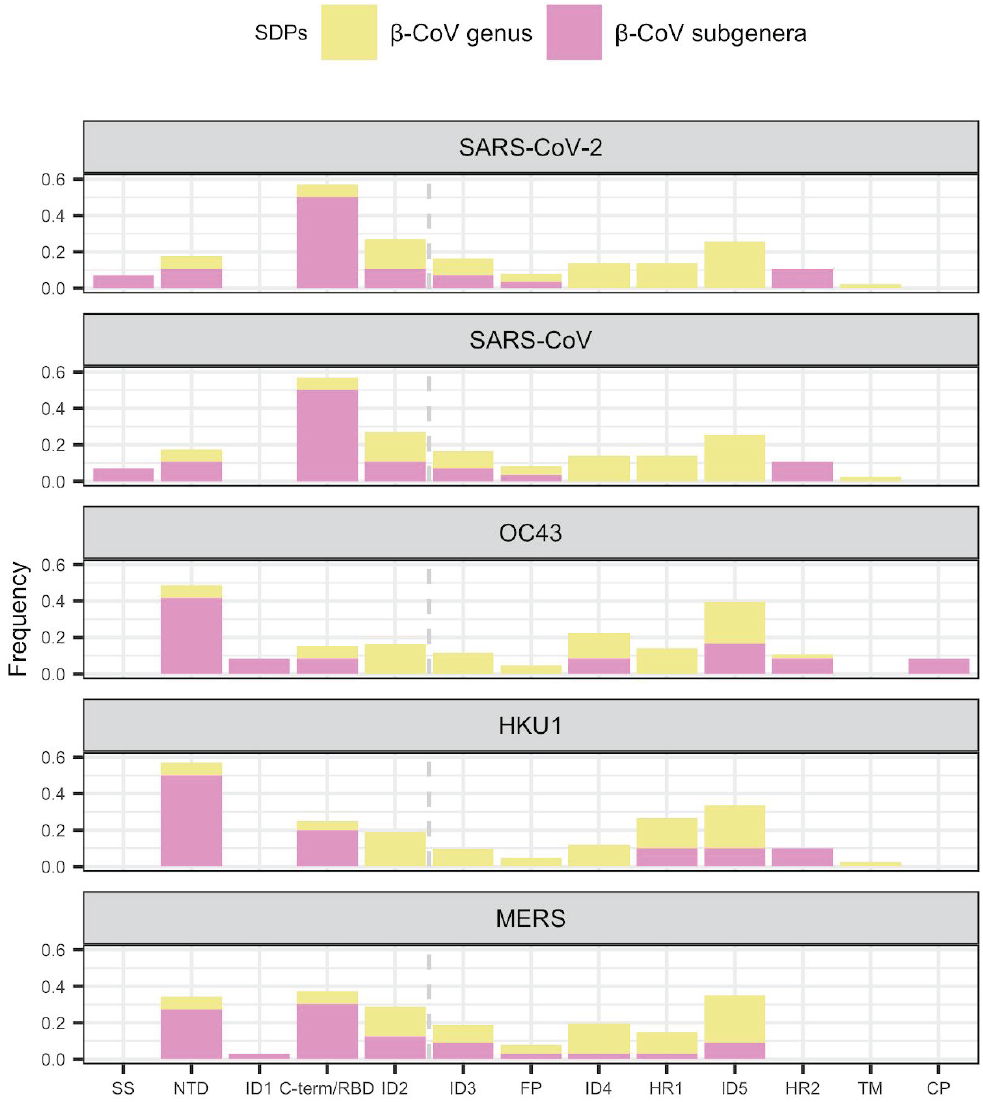
Frequency distribution of SDPs across different domains of the spike sequence from five human pathogenic β-CoVs. Protein domains are denoted as follows: *SS*, signal sequence; *NTD*, N-terminal domain; *RBD*, receptor binding domain; *FP*, fusion peptide; *HR1*, heptad repeat 1; *HR2*, heptad repeat 2. Interdomain regions are denoted by *ID* followed by an integer according to the order in which they appear in the sequence. Dashed vertical lines denote S1/S2 subunits boundaries.

**In summary, the SDPs found within these β-CoV subgenera define a specific region of the receptor binding domains: they are part of, or in direct contact with, the ACE2 interacting surface (Figure 3, Supplementary Figure S3); have a relative low impact on protein stability (Supplementary Figure S4); and have evolved quite recently (Supplementary Figure S5). These characteristics point to a key role of SDPs in mediating the functional specificity of the protein subfamily, i**.**e. the recognition of the host-cell receptor**.

**Figure 3.**
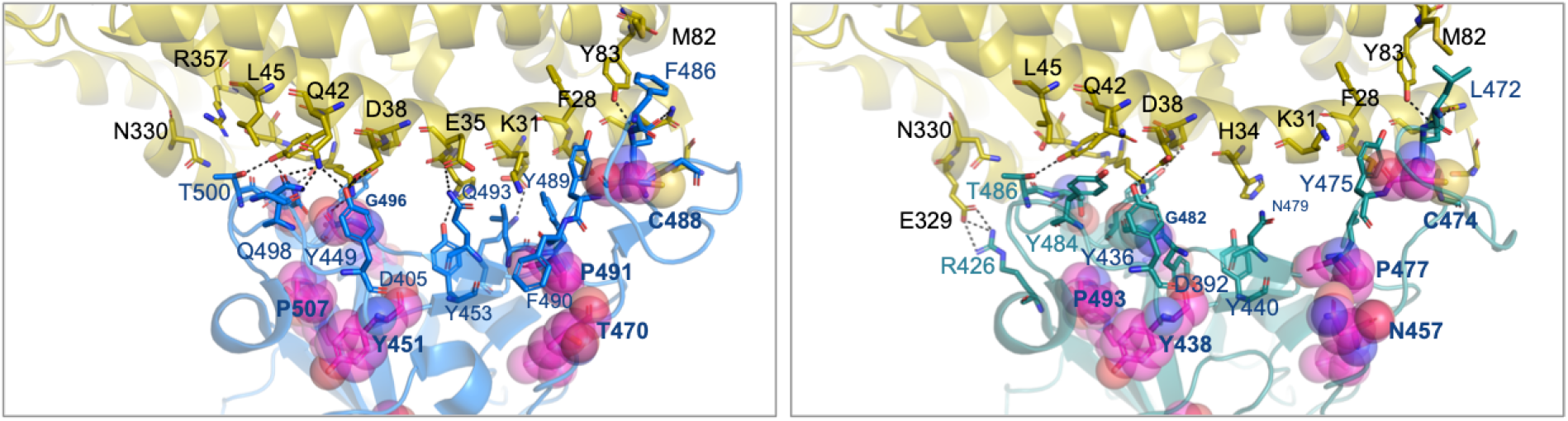
3D structure of the spike protein from SARS-CoV-2 (blue; PDB ID: 6LZG) and SARS-CoV (green; PDB ID: 2AJF) in complex with the human ACE2 cell receptor (yellow). Amino acid residues at the interface are shown as sticks. Intermolecular contacts are shown as dashed black lines. SDPs are highlighted as spheres.

A second S3Det analysis was performed on the full β-CoV MSA. As shown in Figure 1A-D, the identified sequence subfamilies are consistent with the phylogenetic classification of Betacoronavirus into five subgenera (29). In this case, the SDPs linked to the full β-CoV spike family segregation are mainly located in the S2 protein subunit, with a statistically significant enrichment in regions corresponding to interdomain 4, domain HR1 and interdomain 5 (Figure 2, Supplementary Table S1-S2). These SDPs show characteristics of sites under structural constraints (36), *i*.*e*. slowly evolving sites (Supplementary Figure S5) buried within the 3D structure of the protein (Supplementary Figure S3) and potentially destabilizing (see results of *in silico* mutagenesis experiments in Supplementary Figure S4). **Collectively, these results indicate that SDPs capture the functional diversification observed within the individual protein subfamilies, whereby host-cell receptor specificity arises in the context of a structural framework that is specific to each β-CoV phylogenetic group**.

### Relationship between SDPs and ACE2 binding across the Sarbecovirus lineage

Structural and mutagenesis studies have shown that the spike RBD of Sarbecovirus contains all the necessary information for host receptor binding and that a few amino acid substitutions in this region can lead to efficient cross-species transmission (30, 37). Binding to ACE2 is clade-specific and occurs at the carboxy-terminal region of the RBD, by an extended concave loop subdomain which forms the interaction interface with the ACE2 N-terminal helix. Notably, both the ACE2-contacting residues and the surrounding amino acids, collectively referred to as the receptor binding motif (RBM), are required to impart human receptor usage within the Sarbecovirus lineage (30). Consistently, we observe that several SDPs associated with the Sarbecovirus subfamily fall within the RBM, *i*.*e*., Y451/Y438, L461/L448, T470/N457, C488/C474, P491/P477, G496/G482, G502/G488, P507/P493, Y508/Y494 on SARS-CoV-2/SARS-CoV sequences, respectively (Figure 3). Of these, Y494 has been previously reported as critical for ACE2 binding (38). Two SDPs, namely G496 and G502, fall within the receptor interface forming two hydrogen bonds with the ACE2 K353. Other SDPs, such as L461, T470, C488, P491 and G496 make direct contact with ACE2 contacting residues. Hence, these SDPs are likely to play an important role in ACE2 binding by affecting the local orientation of ACE2 contacting residues. This hypothesis is further supported by results from *in silico* mutagenesis and coevolution analysis (details in next section). Specifically, we tested the effect of amino acid mutations across the RBM by mutating the SARS-CoV-2 sequence to the consensus spike sequence from clade 2, i.e. a clade known to be incompatible with hACE2 usage (30). **Notably, while most substitutions are predicted to have a destabilizing effect on the Spike-hACE2 complex, mutations of SDP residues are predicted to have a significantly larger impact compared to non-SDP residues (Figure 4)**.

**Figure 4.**
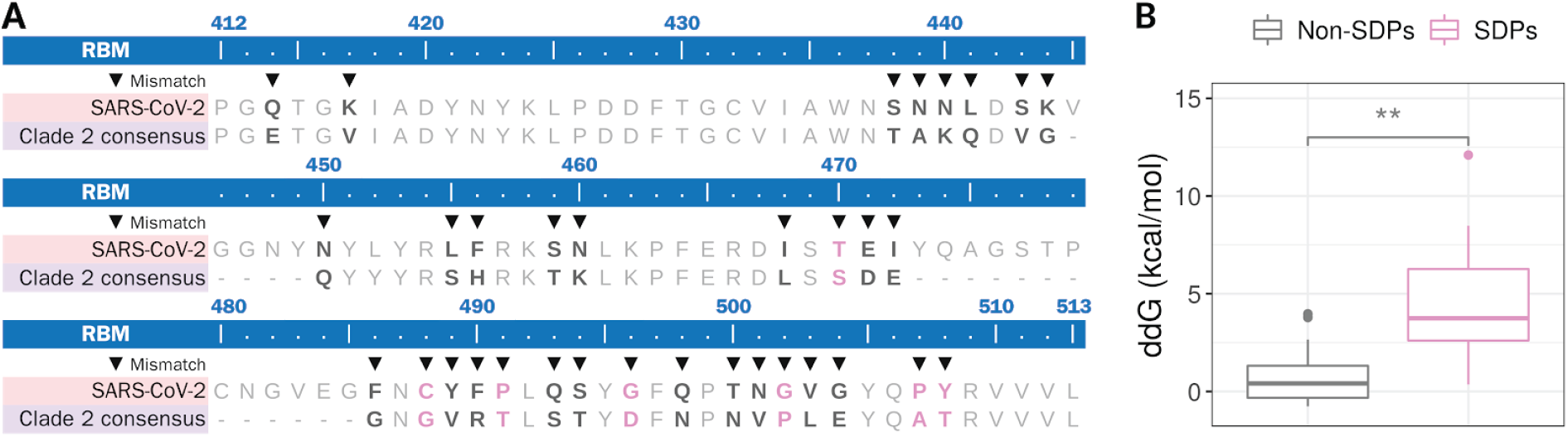
A) Pairwise sequence alignment of SARS-CoV-2 RBM and consensus sequence of Sarbecovirus clade 2. Black triangles indicate amino acid mismatches. SDP positions are depicted as pink letters. B) Boxplot distributions of ΔΔG values resulting from mutating SDPs and non-SDPs using FoldX (PDB ID: 6LZG). Significant differences were computed using a Wilcoxon unpaired two-sample test (p-value < 0.01).

### Molecular coevolution analysis

Computational methods exploiting coevolution signals in MSAs of protein families are widely used to infer features such as molecular interactions and functional sites (15, 39–41). Such signals arise from the specific adaptation between correlated amino acid sites, where changes in one site are potentially compensated by changes in the other. In the case at hand, coevolution signatures are used as markers for the study of the physical interactions occurring between different sites of the spike as well as between the spike and their cognate host cell receptor ACE2. As it can be observed in Supplementary Figure S6, the strongest intramolecular coevolution signal, considering the top-500 predictions, is observed over the RBD region of the spike (the overall precision of the method is reported in Supplementary Figure S7). Figure 5 shows in detail the RBM region, which presents above average precision and recall values of 46% and 4.6%, respectively. Among the SDPs (highlighted in green), the precision is even higher, around 62%, and the recall is 8.7%. **Notably, 18% of SDPs found within the Sarbecovirus subgenus show a coevolution signal with ACE2 contacting residues, namely Y489 (coevolving with C488, P491), Q493 (coevolving with L461, T470, C488, P491) and Q498 (coevolving with G496). These three positions are hubs on the interface with ACE2, making direct contact with positions T27, F28, K31 and Y83, positions K31, H34 and E35, and positions D38, Y41, Q42 and L45 of ACE2, respectively** (22). Particularly, position Q493/N479 in SARS-CoV-2/SARS-CoV has been described to be critical for high affinity binding of both SARS-CoV and SARS-CoV-2 to ACE2 (42, 43). Furthermore, it is interesting to notice that the C488/C474 SDP in SARS-CoV-2/SARS-CoV is an important position for the stability of the RBM as a whole and a complete loss of hACE2 binding *in vitro* has been described when this position is mutated to Alanine in SARS-CoV (44).

**Figure 5.**
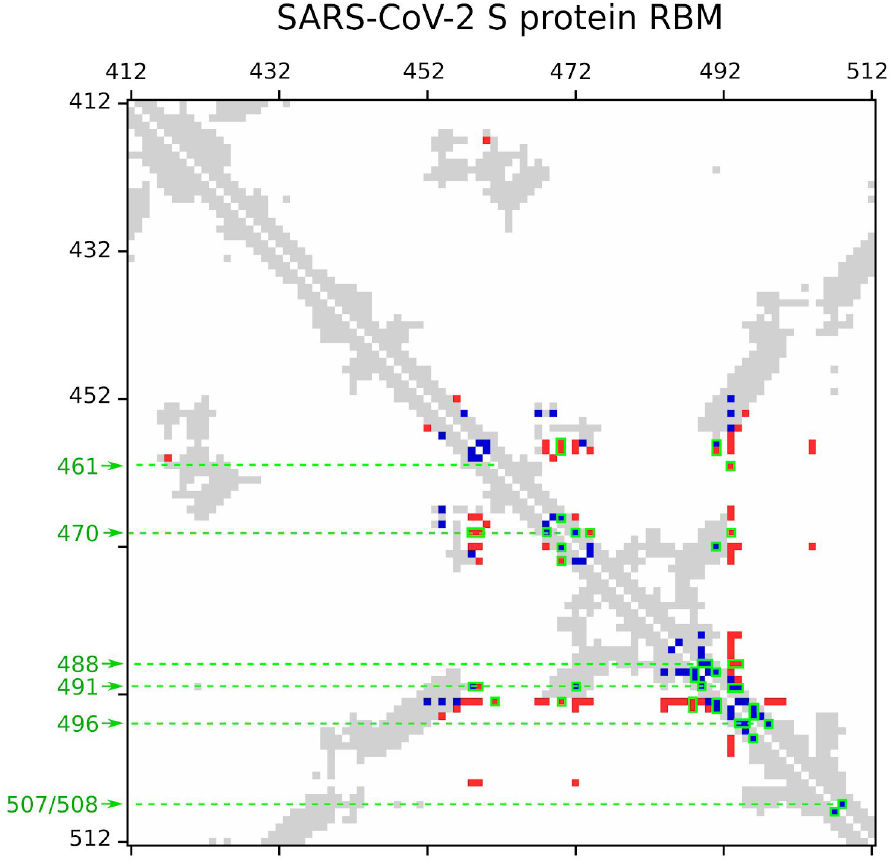
Contact map (8A distance cutoff, any atom) over the RBM of the SARS-CoV-2 spike protein. MI-APC contact predictions (among top 500 scores) are shown in blue (true positives) and in red (false positives). SDPs are highlighted in green.

We next performed a coarse-grained coevolutionary analysis on a concatenated MSA containing eight spike proteins and their cognate ACE2 receptors. Contact predictions were obtained by computing the MI-APC score for every inter-protein pair of alignment positions, considering the RBD region of the spike protein and the whole ACE2. Interestingly, three RBM positions were found coupled to ACE2 positions among the top-10 MI-APC scores (Supplementary Figure S8). Specifically, the ACE2 residue H34 was coupled to L455, S494 and Q498 on the spike protein. Additionally, the spike positions R346 and L455 were coupled to ACE2 residues Q24, N61, Q81, M82 and to E23, T27 and H34, respectively. Among these predictions, T27-L455, H34-L455 and H34-S494 correspond to true contacts within 8A distance, while the couplings between R346 and ACE2 positions could be related to long-range effects. Notably, positions E23 and H34 have been described as crucial to SARS-CoV binding to ACE2 (45). Also, L455 was described as important for the stabilisation of a binding hotspot between SARS-CoV-2 and ACE2 (43). Interestingly, a recent study reported that ACE2 variants E23K and T27A are more susceptible to SARS-CoV-2, while variant H34R decreases SARS-CoV-2 affinity (46). **These results, despite the limited number of sequences in the concatenated alignment, point to at least three specific RBM viral positions (L455, S494 and Q498 in SARS-CoV-2) likely adapted to their species-specific counterparts**.

### Evolution of the SARS-CoV-2 spike protein during the spread in human populations

Since the beginning of the COVID-19 outbreak and the isolation of the SARS-CoV-2 virus, laboratories around the world are continuously isolating viral genomic sequences with unprecedented speed, enabling nearly real-time data sharing of more than 80,000 genomic sequences so far (47).

After discarding partial sequences, a multiple sequence alignment was built based on a total of 44,812 SARS-CoV-2 spike sequences isolated from human samples in 96 countries.

Our analysis of missense amino acid variations confirmed earlier reports (48) that most mutations occur within the S1 subunit, with a dominant variant observed at position 614, where more than 75% of samples carry the D614G mutation, followed by mutations L4F and R21I appearing in 337 and 307 samples, respectively. Within the RBD, the top frequent mutations are S477N, T478I and P479S, found in 68, 63, and 53 samples, respectively. **Notably, these positions fall within the RBM, forming a surface-exposed loop that is proximal to the ACE2 binding surface, and that is absent in Sarbecovirus clades 2 and 3** (30). Although previous experiments indicate that this loop is *per se* not sufficient to impart ACE2 receptor usage, deletions of this region are associated with reduced spike expression and loss of cell entry (30). Hence, mutations in this region are expected to impact the stability of the protein, rather than its affinity to the receptor.

Finally, perhaps not surprisingly, the frequency of variants at SDPs is very low, with an overall variability that is comparable to that observed in ACE2-contacting residues (Supplementary Figure S9). Mutations are observed in 7 out 14 SDPs. With the exception of T385A and Y508, that are observed in 2 and 4 samples, respectively, all other mutations are only present in one sample. Mostly, these variants are predicted to be neutral or to destabilize the binding to the receptor (Supplementary Figure S10B). **In summary, the low frequency of mutations in SDPs fulfils the requirement to preserve a functional role of those positions, providing an additional evidence of their involvement in the interaction with the host cell receptor**.

## Discussion

The relationship between protein family segregation and their functional organisation has been extensively investigated for decades and a variety of computational methods have been developed to infer their evolutionary link at the residue level (49–51). It is therefore relatively straightforward to identify the amino acid positions that modulate the functional specificity of a given enzyme towards a substrate or cofactor (15) or the binding specificity of a protein-ligand or protein-protein interaction (52, 53) by the analysis of the differential conservation patterns within the MSA of a protein family. Here we apply this concept to the analysis of the SARS-CoV-2 spike in the context of a MSA of homologous sequences belonging to the β-CoV genus. The SDP analysis is based on a vectorial representation of protein sequences and amino acid positions in a multidimensional space to simultaneously identify the family segregation and the residue positions that better explain the sources of variation of the family (15).

On one hand, the analysis performed at the β-CoV family level led to the identification of five sequence subfamilies, reflecting the known phylogenetic classification of Betacoronavirus into five subgenera (29) (Figure 1A). On the other hand, the analysis performed on individual β-CoV subgenera, i.e. Sarbecovirus, Merbecovirus and Embecovirus subgroups, allowed a fine-grained classification into subfamily clusters that clearly reflect the functional diversification of the spike protein family, that is, the specificity to different host-cell receptors (Figure 2-3). Indeed, both the clustering and domain enrichment results of the associated SDPs consistently reproduce the known cell receptor specificities observed across the different β-CoV lineages (26, 54–56). At the level of the Sarbecovirus group, for example, both SARS-CoV-2 and RaTG13 are clustered together with SARS-CoV and other SARS-like sequences from bats (Figure 1C), reflecting the ability of these members of the Sarbecovirus group to bind the ACE2 cell receptor (57). Notably, the proximity of SARS-CoV-2 and RaTG13 and other SARS-CoV sequences based on key SDP positions is different to what seen based on full sequence phylogenetic analysis, where they form distinct clades. **A proximity that is driven by their shared SDPs and it is interpreted here in terms of their shared ability to bind the human ACE2 receptor**.

As it is often the case, functional constraints arise from the requirement of maintaining the interaction of proteins with other macromolecules or ligands. Such constraints translate into specific roles and properties of individual amino acids, or protein sites. In the case at hand, the analysis of the physicochemical, structural and conservation properties of the SDPs of the different subfamilies highlights a pattern that is typical of protein functional sites, as they show high conservation across the protein family, are solvent exposed, and are enriched in the receptor binding domains (Supplementary Figures S3-5 and Supplementary Table S2). Hence, in order to assess the role of the SDPs in mediating ACE2 receptor usage across the Sarbecovirus group, we set up an in silico mutagenesis study and analysed the effect of amino acid mutations across the RBM and their impact on ACE2 binding. Notably, while our results are in line with previous observations (30), they point to specific positions across the RBM that might exert a critical role, either by engaging the receptor through direct intermolecular contacts or by affecting the local orientation of ACE2 contacting residues and the stability of the RBM as a whole.

**Collectively, these results point to a key role of SDPs in mediating host cell receptor specificity across β-CoVs and provide, at the same time, a framework for monitoring the evolution of the SARS-CoV-2 specificity to hACE2, as well as the emergence of novel potential cross-species transmission events**. As such, it is important to notice that from the analysis of amino acid variations across the circulating SARS-CoV-2 virus, SDPs tend to mutate with a very low frequency, similar to what is seen at ACE2 contacting sites (Supplementary Figure S9) (58). This is of relevance, as our results suggest that mutations in SDPs can significantly impact the receptor-binding ability of the spike. Furthermore, the experience in other scenarios has shown that mutating SDPs is, in general, sufficient to transform the properties between two groups of proteins of the same family, *i*.*e*. the interchange of the residues occupying the SDPs between two families implies a change in the associated biological properties (59–63). Notable examples include the production of switch-of-function mutants of small GTPases with changed selectivity, or the change of transport specificity between MIP channel proteins by few amino acid substitutions (60, 62). In line with this reasoning, it can be argued that other members of the Sarbecovirus group might have the potential to acquire ACE2 binding ability, as they share substantial similarity in terms of SPDs. This is especially the case of members of the Sarbecovirus green cluster, which despite being phylogenetically distant to SARS-CoV-2, display identical residues in 10 out of 14 SDP positions within the RBD, making them potential candidates for new human infections.

**In conclusion, the results presented here show that the identification of evolutionary patterns based on the analysis of sequence information alone can provide meaningful insights on the molecular basis of host-pathogen interactions and adaptation**. We believe that both the methodology and results presented in this work can provide the basis for follow-up studies analysing the potential routes of mutations that could lead to new adaptation to human hosts and ultimately contribute to better understanding and monitoring of events that are critical to public health concerns worldwide.

## Acknowledgments

The authors are grateful to all other members of the Computational Biology group for their valuable feedbacks and useful discussions.

## References

1. L. S. Hung, The SARS epidemic in Hong Kong: what lessons have we learned? J. R. Soc. Med. 96, 374–378 (2003).

2. F. S. Aleanizy, N. Mohmed, F. Y. Alqahtani, R. A. El Hadi Mohamed, Outbreak of Middle East respiratory syndrome coronavirus in Saudi Arabia: a retrospective study. BMC Infect. Dis. 17, 23 (2017).

3. E. Dong, H. Du, L. Gardner, An interactive web-based dashboard to track COVID-19 in real time. Lancet Infect. Dis. (2020) https://doi.org/10.1016/S1473-3099(20)30120-1.

4. S. Su, et al., Epidemiology, Genetic Recombination, and Pathogenesis of Coronaviruses. Trends Microbiol. 24, 490–502 (2016).

5. S. Xia, et al., Fusion mechanism of 2019-nCoV and fusion inhibitors targeting HR1 domain in spike protein. Cell. Mol. Immunol. (2020) https://doi.org/10.1038/s41423-020-0374-2.

6. F. Li, Evidence for a common evolutionary origin of coronavirus spike protein receptor-binding subunits. J. Virol. 86, 2856–2858 (2012).

7. D. Wrapp, et al., Cryo-EM structure of the 2019-nCoV spike in the prefusion conformation. Science 367, 1260–1263 (2020).

8. J. Lan, et al., Structure of the SARS-CoV-2 spike receptor-binding domain bound to the ACE2 receptor. Nature (2020) https://doi.org/10.1038/s41586-020-2180-5.

9. K. G. Andersen, A. Rambaut, W. Ian Lipkin, E. C. Holmes, R. F. Garry, The proximal origin of SARS-CoV-2. Nat. Med. 26, 450–452 (2020).

10. F. Li, Structure, Function, and Evolution of Coronavirus Spike Proteins. Annu Rev Virol 3, 237–261 (2016).

11. M. F. Boni, et al., Evolutionary origins of the SARS-CoV-2 sarbecovirus lineage responsible for the COVID-19 pandemic. Evolutionary Biology, 83 (2020).

12. S. F. Altschul, W. Gish, W. Miller, E. W. Myers, D. J. Lipman, Basic local alignment search tool. J. Mol. Biol. 215, 403–410 (1990).

13. L. Fu, B. Niu, Z. Zhu, S. Wu, W. Li, CD-HIT: accelerated for clustering the next-generation sequencing data. Bioinformatics 28, 3150–3152 (2012).

14. K. Katoh, J. Rozewicki, K. D. Yamada, MAFFT online service: multiple sequence alignment, interactive sequence choice and visualization. Brief. Bioinform. 20, 1160–1166 (2019).

15. A. Rausell, D. Juan, F. Pazos, A. Valencia, Protein interactions and ligand binding: from protein subfamilies to functional specificity. Proc. Natl. Acad. Sci. U. S. A. 107, 1995–2000 (2010).

16. S. Guindon, et al., New algorithms and methods to estimate maximum-likelihood phylogenies: assessing the performance of PhyML 3.0. Syst. Biol. 59, 307–321 (2010).

17. S. D. Dunn, L. M. Wahl, G. B. Gloor, Mutual information without the influence of phylogeny or entropy dramatically improves residue contact prediction. Bioinformatics 24, 333–340 (2008).

18. S. Xia, et al., Inhibition of SARS-CoV-2 (previously 2019-nCoV) infection by a highly potent pan-coronavirus fusion inhibitor targeting its spike protein that harbors a high capacity to mediate membrane fusion. Cell Res. 30, 343–355 (2020).

19. T. Pupko, R. E. Bell, I. Mayrose, F. Glaser, N. Ben-Tal, Rate4Site: an algorithmic tool for the identification of functional regions in proteins by surface mapping of evolutionary determinants within their homologues. Bioinformatics 18 Suppl 1, S71–7 (2002).

20. D. K. Sydykova, B. R. Jack, S. J. Spielman, C. O. Wilke, Measuring evolutionary rates of proteins in a structural context. F1000Res. 6, 1845 (2017).

21. A. C. Walls, et al., Structure, Function, and Antigenicity of the SARS-CoV-2 Spike Glycoprotein. Cell 181, 281–292.e6 (2020).

22. Q. Wang, et al., Structural and Functional Basis of SARS-CoV-2 Entry by Using Human ACE2. Cell 181, 894–904.e9 (2020).

23. Y. Yuan, et al., Cryo-EM structures of MERS-CoV and SARS-CoV spike glycoproteins reveal the dynamic receptor binding domains. Nat. Commun. 8, 15092 (2017).

24. F. Li, W. Li, M. Farzan, S. C. Harrison, Structure of SARS coronavirus spike receptor-binding domain complexed with receptor. Science 309, 1864–1868 (2005).

25. M. A. Tortorici, et al., Structural basis for human coronavirus attachment to sialic acid receptors. Nat. Struct. Mol. Biol. 26, 481–489 (2019).

26. N. Wang, et al., Structure of MERS-CoV spike receptor-binding domain complexed with human receptor DPP4. Cell Res. 23, 986–993 (2013).

27. R. N. Kirchdoerfer, et al., Pre-fusion structure of a human coronavirus spike protein. Nature 531, 118–121 (2016).

28. S. Guindon, F. Delsuc, J.-F. Dufayard, O. Gascuel, Estimating maximum likelihood phylogenies with PhyML. Methods Mol. Biol. 537, 113–137 (2009).

29. R. Lu, et al., Genomic characterisation and epidemiology of 2019 novel coronavirus: implications for virus origins and receptor binding. Lancet 395, 565–574 (2020).

30. M. Letko, A. Marzi, V. Munster, Functional assessment of cell entry and receptor usage for SARS-CoV-2 and other lineage B betacoronaviruses. Nat Microbiol 5, 562–569 (2020).

31. R. J. G. Hulswit, et al., Human coronaviruses OC43 and HKU1 bind to 9--acetylated sialic acids via a conserved receptor-binding site in spike protein domain A. Proc. Natl. Acad. Sci. U. S. A. 116, 2681–2690 (2019).

32. P. Zhou, et al., A pneumonia outbreak associated with a new coronavirus of probable bat origin. Nature 579, 270–273 (2020).

33. W. Li, et al., Angiotensin-converting enzyme 2 is a functional receptor for the SARS coronavirus. Nature 426, 450–454 (2003).

34. W. Song, et al., Identification of residues on human receptor DPP4 critical for MERS-CoV binding and entry. Virology 471-473, 49–53 (2014).

35. W. Li, et al., Identification of sialic acid-binding function for the Middle East respiratory syndrome coronavirus spike glycoprotein. Proc. Natl. Acad. Sci. U. S. A. 114, E8508–E8517 (2017).

36. A. Tóth-Petróczy, D. S. Tawfik, Slow protein evolutionary rates are dictated by surface-core association. Proc. Natl. Acad. Sci. U. S. A. 108, 11151–11156 (2011).

37. V. D. Menachery, et al., A SARS-like cluster of circulating bat coronaviruses shows potential for human emergence. Nat. Med. 21, 1508–1513 (2015).

38. S. Chakraborti, P. Prabakaran, X. Xiao, D. S. Dimitrov, The SARS Coronavirus S Glycoprotein Receptor Binding Domain: Fine Mapping and Functional Characterization. Virol. J. 2, 1–10 (2005).

39. T. A. Hopf, et al., Sequence co-evolution gives 3D contacts and structures of protein complexes. Elife 3 (2014).

40. D. de Juan, F. Pazos, A. Valencia, Emerging methods in protein co-evolution. Nat. Rev. Genet. 14, 249–261 (2013).

41. J. Rodriguez-Rivas, S. Marsili, D. Juan, A. Valencia, Conservation of coevolving protein interfaces bridges prokaryote-eukaryote homologies in the twilight zone. Proc. Natl. Acad. Sci. U. S. A. 113, 15018–15023 (2016).

42. W. Li, et al., Receptor and viral determinants of SARS-coronavirus adaptation to human ACE2. EMBO J. 24, 1634–1643 (2005).

43. J. Shang, et al., Structural basis of receptor recognition by SARS-CoV-2. Nature (2020) https://doi.org/10.1038/s41586-020-2179-y.

44. S. K. Wong, W. Li, M. J. Moore, H. Choe, M. Farzan, A 193-amino acid fragment of the SARS coronavirus S protein efficiently binds angiotensin-converting enzyme 2. J. Biol. Chem. 279, 3197–3201 (2004).

45. D. P. Han, A. Penn-Nicholson, M. W. Cho, Identification of critical determinants on ACE2 for SARS-CoV entry and development of a potent entry inhibitor. Virology 350, 15–25 (2006).

46. E. W. Stawiski, et al., Human ACE2 receptor polymorphisms predict SARS-CoV-2 susceptibility https://doi.org/10.1101/2020.04.07.024752.

47. S. Elbe, G. Buckland-Merrett, Data, disease and diplomacy: GISAID’s innovative contribution to global health. Global Challenges 1, 33–46 (2017).

48. S. Laha, et al., Characterizations of SARS-CoV-2 mutational profile, spike protein stability and viral transmission. Infect. Genet. Evol. 85, 104445 (2020).

49. A. E. Todd, C. A. Orengo, J. M. Thornton, Evolution of function in protein superfamilies, from a structural perspective. J. Mol. Biol. 307, 1113–1143 (2001).

50. G. Casari, C. Sander, A. Valencia, A method to predict functional residues in proteins. Nat. Struct. Biol. 2, 171–178 (1995).

51. R. A. Laskowski, J. D. Watson, J. M. Thornton, Protein function prediction using local 3D templates. J. Mol. Biol. 351, 614–626 (2005).

52. O. Lichtarge, H. R. Bourne, F. E. Cohen, An evolutionary trace method defines binding surfaces common to protein families. J. Mol. Biol. 257, 342–358 (1996).

53. A. del Sol, F. Pazos, A. Valencia, Automatic methods for predicting functionally important residues. J. Mol. Biol. 326, 1289–1302 (2003).

54. G. Lu, et al., Molecular basis of binding between novel human coronavirus MERS-CoV and its receptor CD26. Nature 500, 227–231 (2013).

55. Q. Wang, et al., Bat origins of MERS-CoV supported by bat coronavirus HKU4 usage of human receptor CD26. Cell Host Microbe 16, 328–337 (2014).

56. Y. Yang, et al., Receptor usage and cell entry of bat coronavirus HKU4 provide insight into bat-to-human transmission of MERS coronavirus. Proc. Natl. Acad. Sci. U. S. A. 111, 12516–12521 (2014).

57. H. Mou, et al., Mutations from bat ACE2 orthologs markedly enhance ACE2-Fc neutralization of SARS-CoV-2 https://doi.org/10.1101/2020.06.29.178459 (August 18, 2020).

58. J. A. Capra, M. Singh, Predicting functionally important residues from sequence conservation. Bioinformatics 23, 1875–1882 (2007).

59. D. Bradley, P. Beltrao, Evolution of protein kinase substrate recognition at the active site. PLoS Biol. 17, e3000341 (2019).

60. V. Lagrée, et al., Switch from an Aquaporin to a Glycerol Channel by Two Amino Acids Substitution. J. Biol. Chem. 274, 6817–6819 (1999).

61. B. Bauer, et al., Effector recognition by the small GTP-binding proteins Ras and Ral. J. Biol. Chem. 274 (1999).

62. W. D. Heo, T. Meyer, Switch-of-Function Mutants Based on Morphology Classification of Ras Superfamily Small GTPases. Cell 113, 315–328 (2003).

63. G. J. Rodriguez, R. Yao, O. Lichtarge, T. G. Wensel, Evolution-guided discovery and recoding of allosteric pathway specificity determinants in psychoactive bioamine receptors. Proc. Natl. Acad. Sci. U. S. A. 107 (2010).

