## Supplementary Materials for "Unraveling the molecular basis of host cell receptor usage in SARS-CoV-2 and other human pathogenic β-CoVs"

#### Supplementary Figures

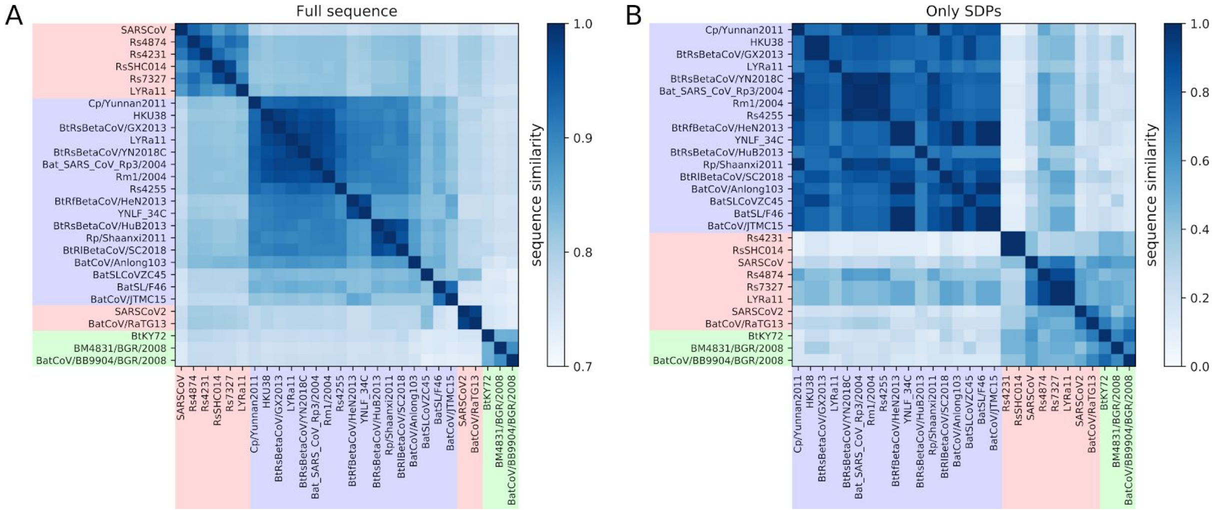


**Figure S1.** All-against-all sequence similarity computed based on Blosum64 substitution matrix on full-length spike protein sequences (A) and only SDPs (B). Sequence labels are colored according to their respective S3Det cluster.

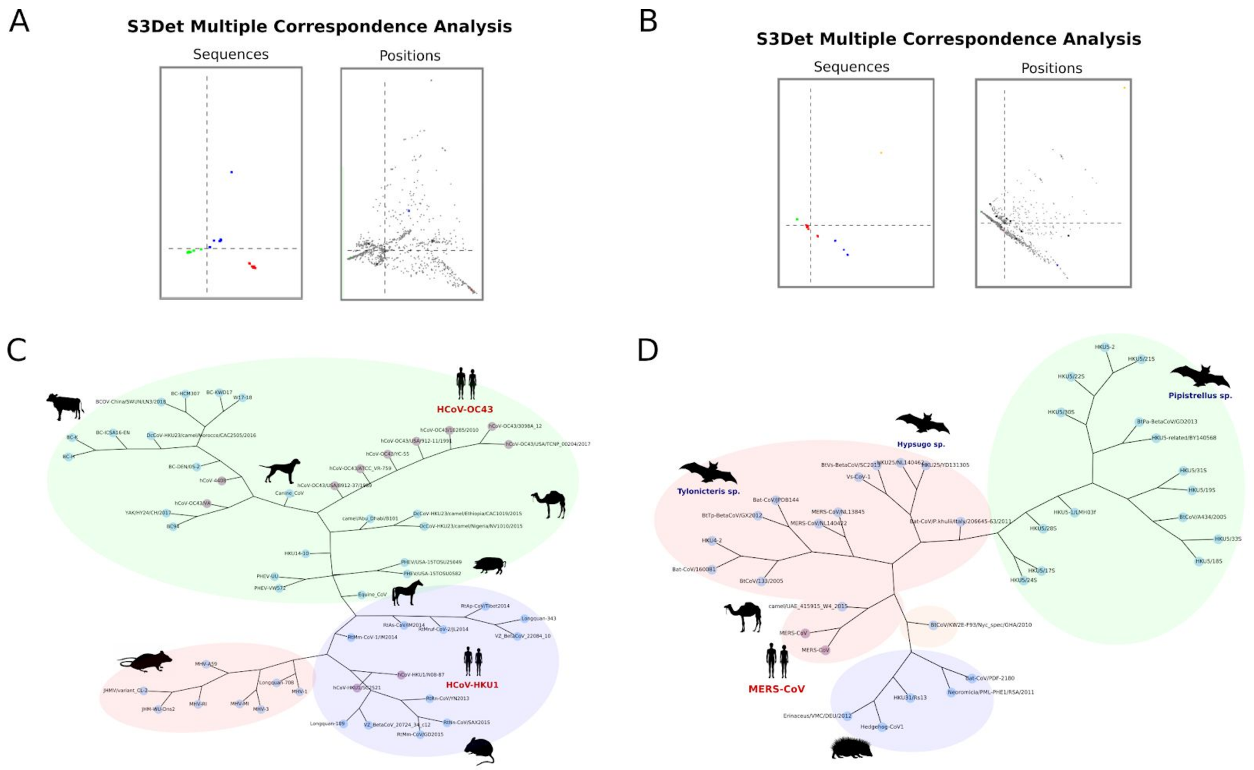


**Figure S2.** Results of the S3Det MCA analysis showing the subfamily segregation and associated amino acid positions obtained for Embecovirus (A) and Merbecovirus (B). Phylogenetic tree of Embecovirus (C) and Merbecovirus (D) spike protein sequences obtained using PhyML[21]. S3Det clusters are highlighted in red, blue, green and orange. Nodes representing the spike protein sequences from human pathogenic CoVs are colored in purple. Host species are shown for some of the nodes as dark silhouettes.

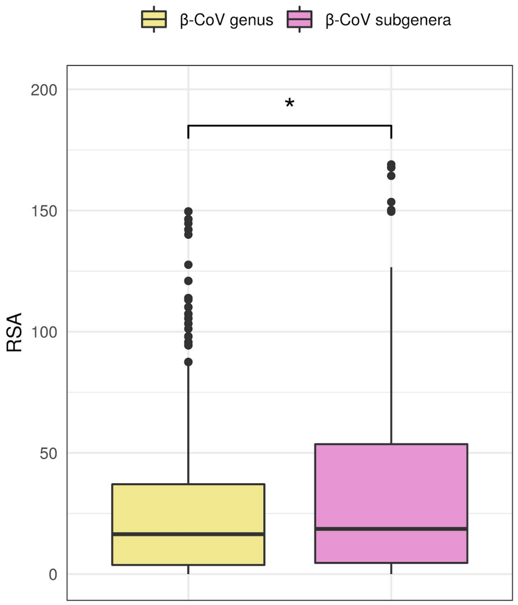


**Figure S3.** Boxplot distribution of per-residue relative solvent accessible area (RSA) for each set of SDPs (β-CoV genus in yellow, β-CoV subgenera in pink). Significant differences were computed using a Wilcoxon unpaired two-sample test (p-value < 0.05).

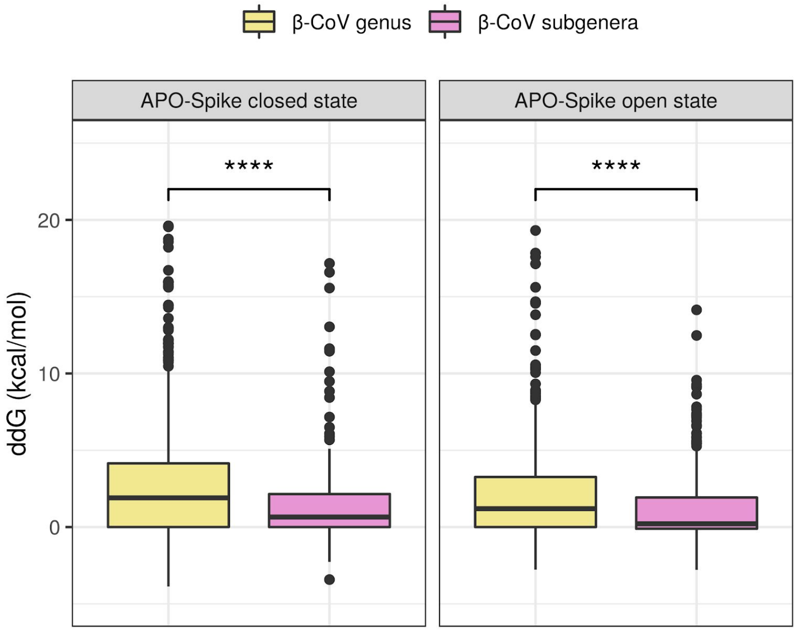


**Figure S4.** Boxplot distribution of ΔΔG values resulting from mutating SDPs associated to the β-CoV genus (yellow) and β-CoV subgenera (pink). ΔΔG values were computed using FoldX (PDB ID: 6VXX, 6VSB). Significant differences were computed using a Wilcoxon unpaired two-sample test (p-value < 0.0001).

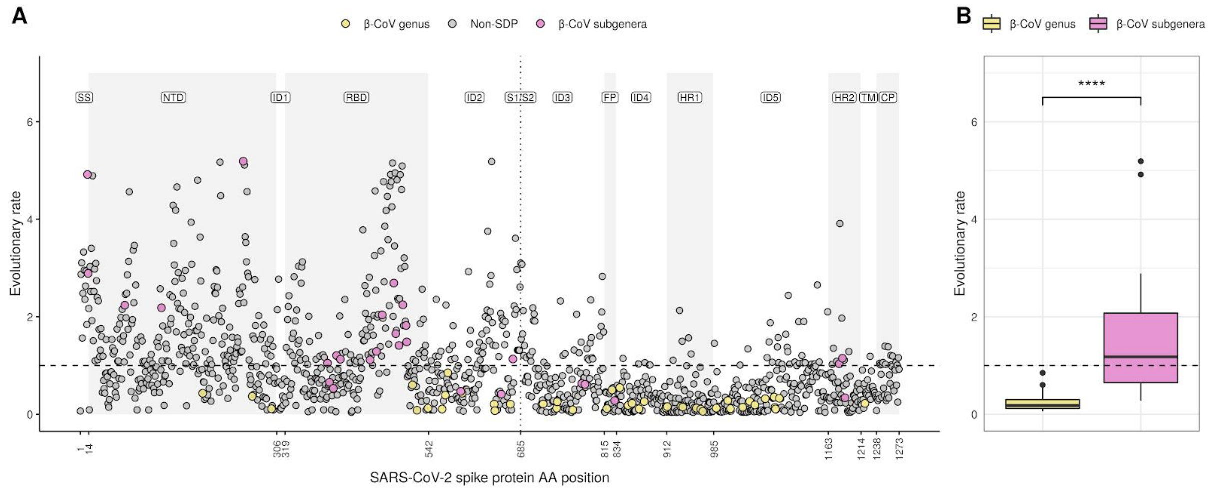


**Figure S5. Evolutionary rate analysis. A)** Site-specific evolutionary rate of the SARS-CoV-2 spike protein as computed by Rate4Site. The dashed horizontal line indicates the mean EV rate. Yellow and pink dots indicate SDPs linked to the β-CoV genus and subgenera, respectively. Protein domains are highlighted by a grey shade and denoted as follows: *SS,* signal sequence; *NTD,* N-terminal domain; *RBD,* receptor binding domain; *FP,* fusion peptide; *HR1,* heptad repeat 1; *HR2*, heptad repeat 2; TM, transmembrane region; CP, cytoplasmic. Interdomain regions are denoted by ID followed by an integer according to the order in which they appear in the sequence*.* The dotted vertical line denotes the S1/S2 subunits boundary. **B)** Boxplot distributions of site-specific evolutionary rates computed on the two sets of SDPs. Significant differences were computed using a Wilcoxon unpaired two-sample test (p-value < 0.0001).

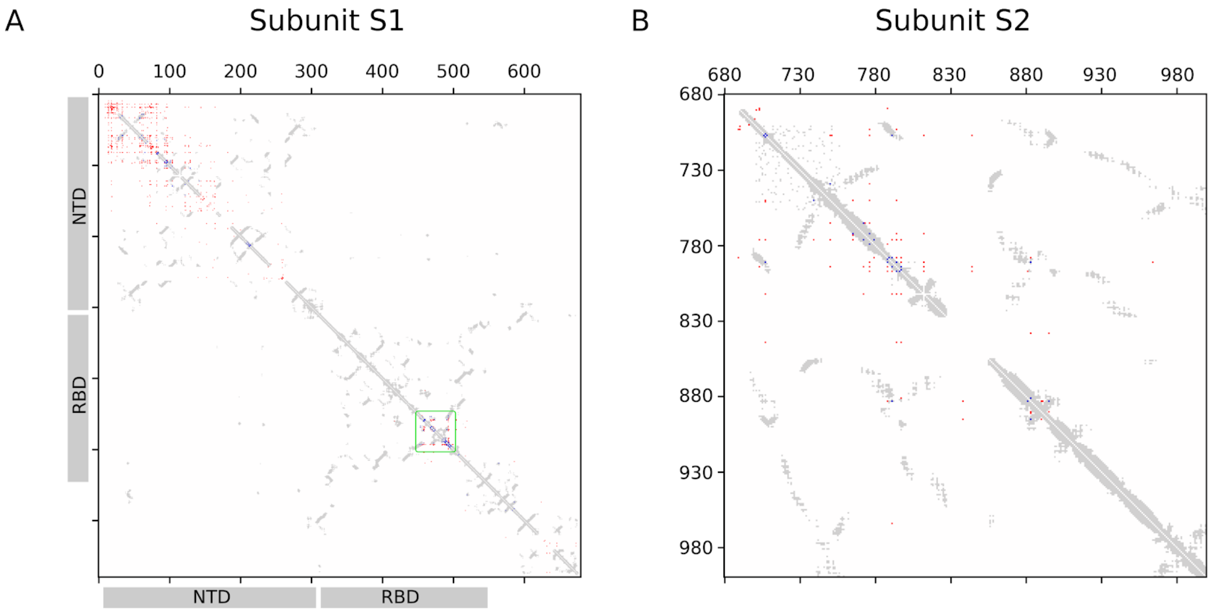


**Figure S6.** Contact map (8A distance cutoff, any atom) for subunit S1 (A) and S2 (B) of the SARS-CoV-2 spike protein. Top-500 MI-APC contact predictions are shown in blue (true positives) and in red (false positives). The RBM is highlighted in green.

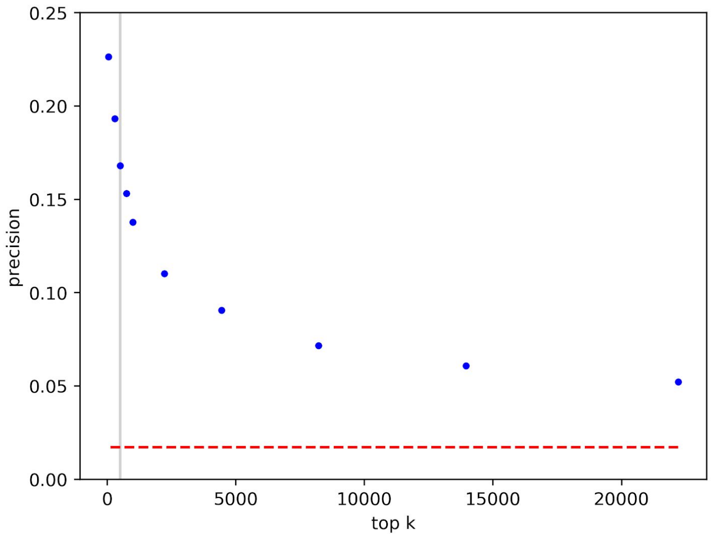


**Figure S7.** Probability of finding a true contact (8A distance cutoff, any atom) among the top-k intra-protein MI-APC predictions for the spike protein. Gray line: top-500, precision = 16.8%. Red dashed line: null model precision = 1.7%.

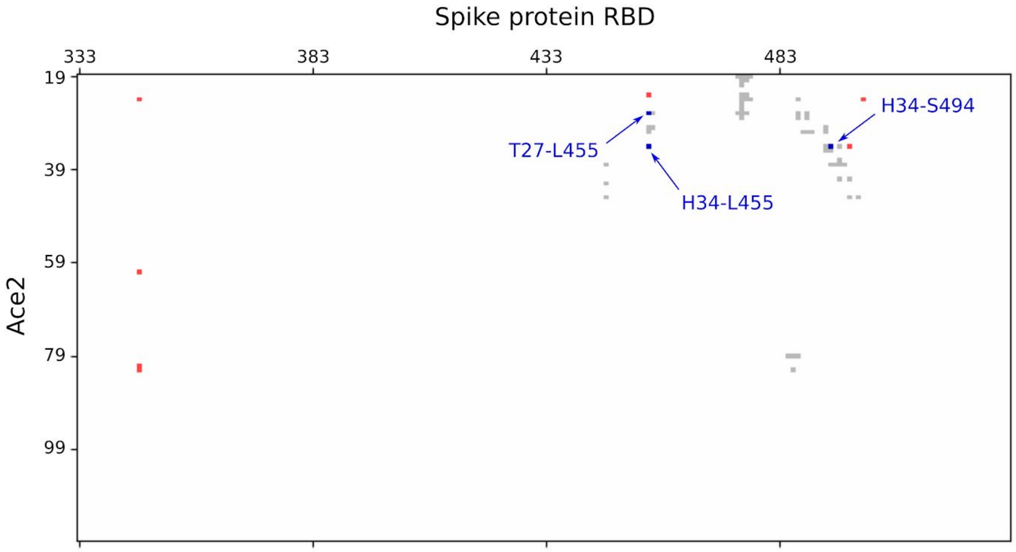


**Figure S8.** Contact map (8A distance cutoff, any atom) between SARS-CoV-2 spike protein RBD and its receptor ace2. The top-10 MI-APC contact predictions are shown in blue (true contacts) and in red (false positives). The contact map was obtained from PDB structure 6LZG.

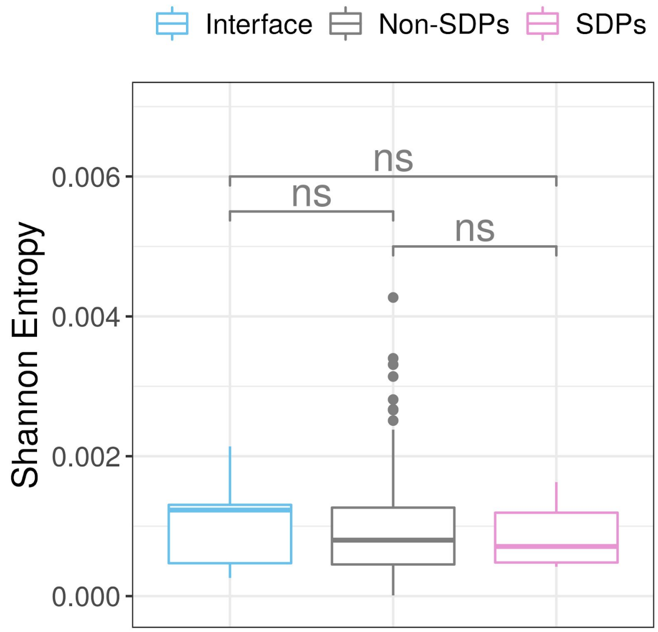


**Figure S9.** Boxplot depicting the level of conservation (Shannon Entropy) computed for SDPs or non-SDPs residues belonging to the RBD.

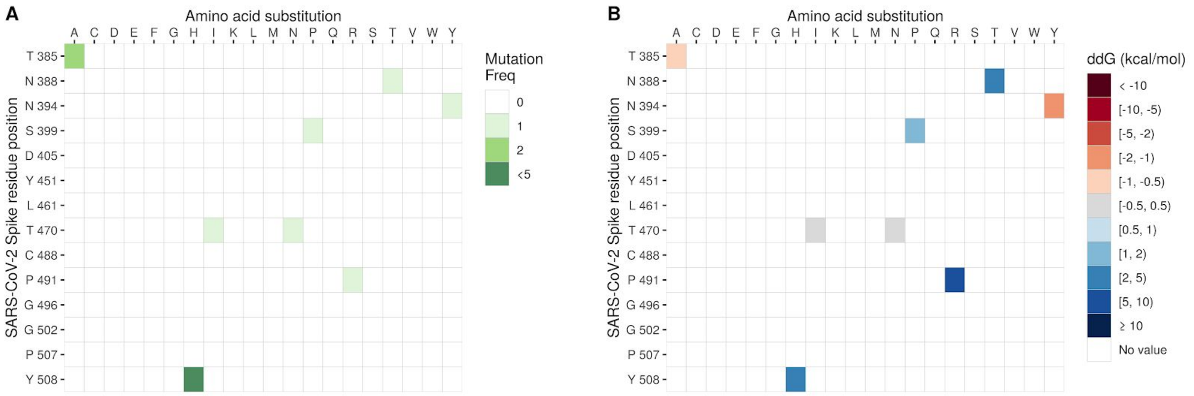


**Figure S10**. A) Frequency of amino acid variations at SDP positions within the RBD. B) Predicted effect of mutations in terms of ΔΔG values computed by Foldx (PDB code: 6LZG).

### Supplementary tables

**Table S1. Subunit-level enrichment analysis.**

| | | p-value hypergeometric test | | | | |
|---|---|---|---|---|---|---|
| **Subunit** | **SDP** | **SARS-CoV-2** | **SARS-CoV** | **OC43** | **HKU1** | **MERS** |
| S1 | β-CoV genus | 0.999574 | 0.999418 | 0.999871 | 0.999742 | 0.999698 |
| | β-CoV subgenera | **0.005667** | **0.004632** | 0.561269 | 0.284334 | **0.031552** |
| S2 | β-CoV genus | **0.001289** | **0.001716** | **0.000429** | **0.000819** | **0.000947** |
| | β-CoV subgenera | 0.998436 | 0.998753 | 0.665795 | 0.891131 | 0.987258 |

Numbers in bold indicate p-value < 0.05.

**Table S2. Domain-level enrichment analysis.**

| | | p-value hypergeometric test | | | | |
|---|---|---|---|---|---|---|
| **Domain** | **SDP** | **SARS-CoV-2** | **SARS-CoV** | **OC43** | **HKU1** | **MERS** |
| SS | β-CoV genus | - | - | - | - | - |
| | β-CoV subgenera | **0.031339** | **0.04214** | - | - | - |
| NTD | β-CoV genus | 0.998889 | 0.998348 | 0.997644 | 0.995925 | 0.999234 |
| | β-CoV subgenera | 0.972201 | 0.965056 | **0.090286** | **0.03567** | 0.406908 |
| ID1 | β-CoV genus | - | - | - | - | - |
| | β-CoV subgenera | - | - | 0.109207 | - | 0.276512 |
| C-Term/RBD | β-CoV genus | 0.988181 | 0.988995 | 0.998751 | 0.999818 | 0.987616 |
| | β-CoV subgenera | **0.000069** | **0.00077** | 0.95516 | 0.713181 | **0.053881** |
| ID2 | β-CoV genus | 0.204935 | 0.195065 | 0.118511 | 0.047175 | 0.137046 |
| | β-CoV subgenera | 0.630695 | 0.620801 | - | - | 0.453568 |
| ID3 | β-CoV genus | 0.649939 | 0.661564 | 0.49089 | 0.671802 | 0.650804 |
| | β-CoV subgenera | 0.794561 | 0.801416 | - | - | 0.683688 |
| FP | β-CoV genus | 0.132685 | 0.135816 | 0.118811 | 0.115631 | 0.116061 |
| | β-CoV subgenera | 0.346639 | 0.35069 | - | - | 0.377577 |
| ID4 | β-CoV genus | **0.043054** | **0.04567** | **0.025754** | **0.075013** | **0.009901** |
| | β-CoV subgenera | - | - | 0.490168 | - | 0.870504 |
| HR1 | β-CoV genus | **0.032391** | **0.034422** | **0.024234** | **0.005911** | 0.072107 |
| | β-CoV subgenera | - | - | - | 0.427303 | 0.844344 |
| ID5 | β-CoV genus | **0.028881** | **0.031754** | **0.046297** | **0.04176** | **0.023204** |
| | β-CoV subgenera | - | - | 0.481971 | 0.759839 | 0.858372 |
| HR2 | β-CoV genus | - | - | 0.805111 | - | - |
| | β-CoV subgenera | 0.098164 | 0.101424 | 0.363018 | 0.320312 | - |
| TM | β-CoV genus | 0.564982 | 0.570284 | - | 0.534476 | - |
| | β-CoV subgenera | - | - | - | - | - |
| CP | β-CoV genus | - | - | - | - | - |
| | β-CoV subgenera | - | - | 0.249255 | - | - |

Numbers in bold indicate p-value < 0.05.
